# Beta burst characteristics and coupling within the sensorimotor cortical-subthalamic nucleus circuit in Parkinson’s disease

**DOI:** 10.1101/2024.11.19.624397

**Authors:** Pan Yao, Bahman Abdi-Sargezeh, Abhinav Sharma, Tao Liu, Huiling Tan, Amelia Hahn, Philip Starr, Simon Little, Ashwini Oswal

## Abstract

**Background:** Bursts of exaggerated subthalamic nucleus (STN) beta activity contribute to clinical impairments in Parkinson’s disease (PD). Few studies have explored the characteristics and coupling of bursts across the sensorimotor cortical-STN circuit.

**Objective:** We sought to (1) establish the characteristics of sensorimotor cortical and STN bursts during naturalistic behaviours, and (2) determine the predictability of STN bursts from motor cortical recordings.

**Methods:** We analysed 1,478 hours of wirelessly streamed bilateral sensorimotor cortical and STN recordings from 5 PD patients.

**Results:** STN bursts were longer than cortical bursts and had shorter inter-burst intervals. Long bursts (>200ms) in both structures displayed temporal overlap (>30%), with an estimated cortico-STN conduction delay of 8ms. Furthermore, approximately 27% of all STN bursts were preceded by a cortical burst.

**Conclusion:** Cortical beta bursts tend to precede STN beta bursts, with short delays. However, subcortical mechanisms are also likely to contribute to STN burst initiation and propagation.

## Introduction

Exaggerated basal ganglia beta (13-30 Hz) frequency oscillatory synchronisation is a hallmark of Parkinson’s disease (PD)^1–3^. Beta activity is not continuous, often occurring in transient packets known as bursts^4,5^. Previous work has demonstrated that beta bursts of prolonged duration and amplitude correlate with motoric impairments^6,7^. Furthermore, therapies such as STN deep brain stimulation (DBS) and dopaminergic medication lead to a reduction in STN beta activity, with the extent of suppression correlating with clinical improvements ^8,9^. These observations have led to beta activity being used as a control signal in amplitude responsive adaptive DBS (aDBS), where STN stimulation is delivered only following the detection of beta bursts and not continuously^10–13^. Growing evidence suggests that aDBS may be more efficacious than continuous DBS^13^.

Although the pathophysiological significance of beta bursts is established, we still do not fully understand the mechanisms leading to burst initiation. Studies of simultaneous basal ganglia and sensorimotor cortical recordings highlight the existence of coherent beta networks in which sensorimotor cortical beta activity drives basal ganglia activity over the same frequency range ^8,14–16^. This finding is consistent with canonical corticobasal ganglia circuitry^17^ and leads to the hypothesis that beta bursts originating within sensorimotor cortex could drive the onset of basal ganglia bursts, therefore reliably preceding their onset. Addressing this question is important, as it may imply predictability of the timing of basal ganglia bursts from sensorimotor cortical recordings, providing knowledge that could be used in aDBS applications or the development of less invasive strategies for modulating deep brain activity based on cortical stimulation.

Here we leverage an investigational DBS device capable of performing simultaneous invasive sensorimotor cortical and STN recordings in a home environment to: (1) establish the characteristics of motor cortical and STN bursts during naturalistic behaviours, and (2) determine the predictability of STN bursts from cortical activity.

## Methods

### Patient characteristics

We analysed data from five patients with PD (see **Supplementary Table 1**), with bilateral implants of the Summit RC+S neural interface (Medtronic). Each STN was implanted with a quadripolar Medtronic 3389 lead, whilst cortical recordings were performed using a quadripolar paddle-type lead (10mm intercontact spacing), such that 2-3 contacts were anterior to the central sulcus, 2-4 cm lateral from the midline. STN and cortical contact locations were confirmed by fusion of post-operative CT and pre-operative MRI imaging as previously reported^18,19^.

Data from two bipolar STN channels and two bipolar cortical channels was streamed from each hemisphere (sampling rate of 250 Hz) to a Microsoft Windows tablet^19^. Recordings were completed between 2-4 weeks after implantation, during activities of daily living, whilst patients were on their usual dopaminergic medication, before the initiation of DBS.

### Defining burst characteristics and the overlap of cortico-STN bursts

Data from each STN and cortical channel was bandpass filtered (±3 Hz using a 4^th^ order zero-phase Butterworth filter) around the beta peak frequency for cortico-STN coherence for each hemisphere (see **Supplementary Table 1** for further details). This allowed us consider bursts at the same frequency that were synchronized across sites. Beta burst timings were defined separately for each channel as time points where the beta amplitude envelope exceeded its 75^th^ percentile^4^. Burst duration and inter-burst interval probability density functions were then computed using 20ms spaced bins for values ranging from 0-1000ms. A GLM (using spm_ancova.m^20^, with covariates added to account for between and within (hemispheric) subject differences) enabled comparison of cortical and STN burst durations and inter-burst intervals at each time bin, with cluster-based permutation testing being used to correct for multiple comparisons (cluster forming and cluster extent thresholds set to P<0.01).

Bursts were also separately defined for each integer frequency (±3 Hz window) ranging from 10 to 35 Hz. For each frequency and for each of the four combinations of STN and cortical channels from each hemisphere, we computed the overlap of STN and cortical bursts. This was defined as the total number of timepoints during which there was a burst in both sites, divided by the number of timepoints during which an STN burst occurred. Overlap proportions were computed separately for short (<200 ms) and long (>200 ms) duration bursts, as only the latter are believed to adopt a pathological role^4,5,21^.

### Generation of surrogate data

Surrogate data allowed us to determine whether observed cortico-STN burst overlap profiles were greater than above chance for both long and short duration bursts. Based on the probability of each 20ms wide burst duration bin (**Figure 1a**) a fixed number of bursts for each bin duration was generated and used to create binarised burst timing data that preserved the original burst characteristics for both sites. For each instantiation of surrogate data for both STN and cortical sites, the ordering of bursts was randomised, and inter-burst intervals were drawn from corresponding density functions (**Figure 1a)**. 1000 instantiations of surrogate data were used to compute null distributions for statistical testing.

**Figure 1.**
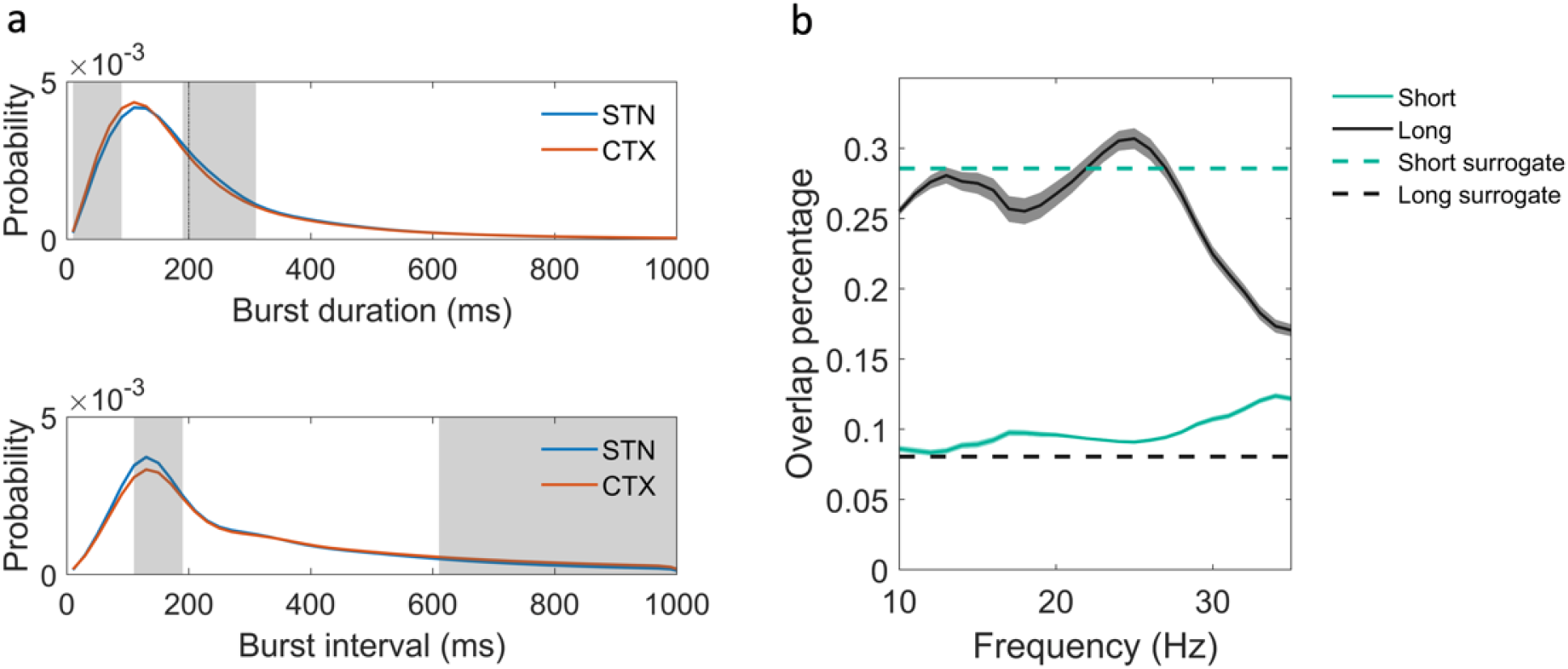
Beta burst characteristics and coupling within the motor cortical-STN circuit. (a) shows probability density functions (pdfs) for STN and motor cortical burst durations and inter-burst intervals. The dotted black line in the burst duration pdf indicates the 200ms cut off that was used to differentiate long and short bursts. There was a higher proportion of short duration bursts (10-90ms) in the cortex and a correspondingly higher proportion of longer duration bursts (190-310ms) within the STN, as indicated by the grey shaded regions. Similarly, the probability of inter-burst intervals between 110-190ms was greater in the STN, whilst the probability of intervals greater than 610ms was greater in the motor cortex. (b) displays the overlap of cortical bursts relative to STN bursts for frequencies in the range of 10-35 Hz (1 Hz resolution ± 3 Hz), separately for long (>200ms) and short (<200ms) burst durations. For long bursts, the maximal overlap was greater than 0.3 and occurred at a frequency of 25Hz. The dashed lines indicate the 99th percentiles of overlaps derived from surrogate data that were generated using the pdfs of observed data plotted in (a). Surrogate data overlaps were computed separately for long and short bursts.

### Directionality of cortico-STN burst coupling

To determine whether long cortical bursts tend to precede long STN bursts, we computed the burst overlap proportion at different time lags for STN timeseries shifts relative to cortical activity. This was performed for the frequency displaying the largest cortico-STN burst overlap. A one-sample t-test was then used to test for statistical significance. Additionally, for both sites, we also computed the proportion of long bursts in one site whose onset was accompanied by and also preceded by a burst in the other site. This was performed separately for both the real and surrogate datasets. Finally, in view of the overlap being maximal at a lag of −8ms (see **Figure 2a**), we computed the proportion of long bursts in one site that were preceded by the onset of a burst in the other site within time windows of 4, 8, 12 and 16ms.

**Figure 2.**
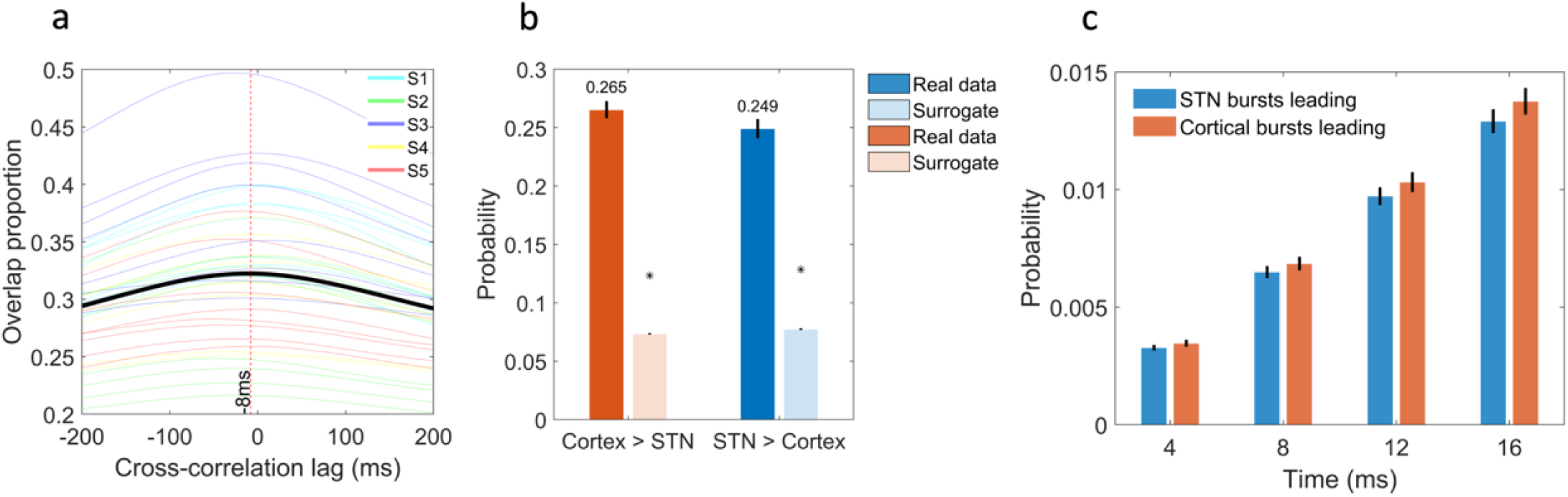
Cortical bursts at the peak overlap frequency tend to precede STN bursts. (a) shows the cross-correlation function of the overlap of long cortical bursts relative to STN bursts at a frequency of 25 ± 3 Hz. Data is shown for individual combinations of STN and cortical channels (4 combinations from 2 bipolar STN and 2 bipolar cortical channels) for each hemisphere of each subject (see legend), with the black line indicating the mean. The maximal overlap is achieved when STN data are shifted −8ms relative to cortical data – indicative of a cortico-STN burst conduction time of 8ms. (b) reveals the proportion of long STN bursts (at a centre frequency of 25 Hz) that were preceded by a cortical burst and vice versa for both the real data and surrogate data. The bars indicate mean values, whilst black lines indicate standard errors. The stars indicate the 99th percentile derived from surrogate data distributions. The onset of 26.5% of all STN bursts was preceded by the onset of a cortical burst, whilst the corresponding figure for STN bursts preceding cortical bursts was 24.9%. Both values were above chance (p<0.01). (c) The proportion of STN bursts (at a centre frequency of 25 Hz) whose onset was preceded by the onset of a cortical burst (and vice versa) within different timeframes (4,8,12 and 16ms) is shown. Bars indicate mean values, whilst the black lines represent standard errors. A 2-factor ANOVA revealed that STN bursts are more likely to be preceded by cortical bursts than vice versa (T_303_ = 2.47, p = 0.007; see Results).

## Results

### Sensorimotor cortical and STN beta burst characteristics

Probability density functions for cortical and STN burst durations, and inter-burst intervals are displayed in **Figure 1a**. Grey regions indicate the presence of significant differences between the STN and cortex. We observed a greater proportion of short duration (10-90 ms) bursts within the sensorimotor cortex and a correspondingly higher proportion of long duration (190-310 ms) bursts within the STN. Furthermore, inter-burst intervals tended to be shorter within the STN than within the sensorimotor cortex.

### Cortico-STN burst overlap and estimation of cortico-STN transmission delays

Long bursts (>200ms) displayed a greater overlap between the sensorimotor cortex and STN than short bursts (<200ms), such that the maximal overlap for long bursts was greater than 30% at the high beta frequency of 25 Hz (**Figure 1b**). Overlap proportions were also derived for surrogate data, with **Figure 1b** showing the 99^th^ percentile of the surrogate distribution for long and short burst overlaps. Only long bursts displayed above chance coupling at all frequencies between 10-35 Hz (significant when corrected for multiple comparisons using cluster-based permutation testing with cluster forming and cluster extent thresholds of P <0.01).

Cross-correlation of cortical and STN burst time courses at the peak overlap frequency of 25 Hz revealed a maximal overlap at mean lag of −8ms (one sample t-test: T_30_ = 8.16, p < 0.01) – suggesting that that cortical bursts tend to precede STN bursts by ∽8ms (**Figure 2a**).

### Cortical bursts precede STN bursts

**Figure 2b** highlights that at the centre frequency of 25 Hz, 26.5% of long STN bursts were preceded by the onset of a sensorimotor cortical burst. Interestingly, we also observed a reciprocal relationship, such that ∽25% of long cortical bursts were preceded by the onset of an STN burst. Both of these percentages were above chance values (P<0.01 indicated by the stars in **Figure 2b**) derived from surrogate data. This result highlights that long bursts in the STN can be preceded by a burst in the cortex, but also vice versa.

Next, we tested whether cortical bursts were more likely to precede long STN bursts than STN bursts preceding long cortical bursts. Given that cross-correlation lags resulting in maximal overlap tended to be in the range of 4-16ms (see **Figure 2a**), we explored whether bursts in one site were preceded by a burst within the other site for values within this time frame. The results, shown in **Figure 2c**, highlight that cortical bursts are more likely to precede STN bursts than vice versa. This was confirmed using a 2-factor ANOVA, with factors leading site (levels: STN leading vs. cortex leading) and precession timeframe (levels: 4ms vs. 8ms vs. 12ms vs. 16ms). There was a main effect of leading site (T_303_ = 2.47, p = 0.007) such that the cortex was more likely to lead than the STN.

## Discussion

We leveraged prolonged invasive recordings of sensorimotor cortical and STN activity during daily living, to interrogate the characteristics of cortical and STN beta bursts. To our knowledge this is the first study to reveal differences in the properties of motor cortical and STN beta bursts in PD– with STN bursts tending to be longer and having shorter inter-burst intervals. Additionally, our findings imply significant cross-site coupling of long bursts, with an overlap percentage of >30% at high beta (25 Hz) frequencies. Notably, this overlap value is markedly greater than that reported in previous studies, where cortical activity has been recorded (with lower signal-to-noise ratio) using EEG^22^. Nevertheless, our results also reveal that STN bursts most commonly occur in the absence of a concurrent cortical burst, a finding which may suggest a role for subcortical pathways in STN burst propagation (e.g., striatal pathways or the reciprocal STN-GPe loop^23–25^).

Cross-correlation analyses revealed that long cortical bursts tend to preced the onset of long STN bursts, with an estimated conduction delay of 8ms. This value is consistent with reports of monosynaptic hyperdirect pathway latencies between the sensorimotor cortex and the STN^26^. Interestingly, we also observed that STN bursts were more likely than chance to precede cortical bursts. This is perhaps unsurprising given the presence of reciprocal anatomical connectivity between the STN and sensorimotor cortex via thalamo-cortical circuits^27^.

Taken together, our results imply that long beta bursts are coupled across the sensorimotor cortical-STN circuit, with cortical bursts tending to precede the onset of STN bursts. These findings also lead us to speculate that suppressing cortical bursts, via non-invasive or minimally invasive approaches, could lead to the suppression of STN bursts and therefore to improvements in motor symptoms.

## Acknowledgements

PY acknowledges support from the China Scholarship Council Program (Project ID 202304910538). This work was supported by an MRC Clinician Scientist Fellowship (MR/W024810/1) held by AO. BA and AO acknowledge funding support from the Oxford University Hospitals Charity. HT is supported by the Medical Research Council UK (MC_UU_00003/2, MR/V00655X/1, MR/P012272/1), the National Institute for Health Research (NIHR) Oxford Biomedical Research Centre (BRC) and the Rosetrees Trust, UK.

## Author roles

(1) Research project: A. Conception, B. Organization, C. Execution; (2) Statistical analysis: A. Design, B. Execution, C. Review and critique; (3) Manuscript preparation: A. Writing of the first draft, B. Review and critique.

P.Y.: 1ABC,2ABC,3AB B.A.:1BC,2C,3B

A.S.:1BC,2C,3B

T.L..:1C,2C,3B

H.T..:1C,2C,3B

A.H..:1C,2C,3B

P.S.: 1ABC,2C,3B

S.L..:1ABC,2C,3B A.O.: 1ABC,2ABC,3B

## Data Availability Statement

Fully anonymized data will be shared with qualified investigators upon request to the corresponding authors.

