## Supplementary Table 1 for "Beta burst characteristics and coupling within the sensorimotor cortical-subthalamic nucleus circuit in Parkinson’s disease"

**Supplementary Materials**

**Supplementary Table 1**

| **Patient** | **Age and gender** | **Disease duration (years)** | | **Preoperative medication (mg)** | **UPDRS III off medication** | **% Change in UPDRS III when on** | **MOCA** | **Data duration for right and left hemispheres**  **(hours)** | **Beta peak frequency for cortico-STN coherence** |
| --- | --- | --- | --- | --- | --- | --- | --- | --- | --- |
| 1 02 | 54 M | | 7 | LDE 1425 | 49 | 90% | 26 | L: 96.9  R: 121.2 | L: 22 Hz  R: 26 Hz |
| 2 05 | 63 M | | 19 | LDE 955 | 45 | 51% | 30 | L: 152.9  R: 153.3 | L: 30 Hz  R: 30 Hz |
| 3 06 | 28 F | | 12 | LDE 1550 | 61 | 73% | 27 | L: 88.6  R: 78.2 | L: 26 Hz  R:28 Hz |
| 4 07 | 40 M | | 4 | LDE 1314 | 41 | 65% | 30 | L: 285.7  R: 274.9 | L: 25 Hz  R: 25 Hz |
| 5 08 | 58 M | | 12 | LDE 2100 | 44 | 75% | 27 | L: 39.4  R: 187.4 | L: 24 Hz  R: 24 Hz |

**Clinical characteristics of patients.** LDE = levodopa dose equivalent. The total pre-operative UPDRS part III score is presented in the off-medication state. The beta peak frequency for cortico-STN coherence was calculated as the peak value within the frequency range (13-30Hz). Coherence was calculated using a multitaper approach (implemented in the FieldTrip toolbox; <https://www.fieldtriptoolbox.org/>), with a frequency resolution of 1Hz and a taper smoothing frequency of 2 Hz.
